# High-Resolution Subtyping of Pediatric Low-Grade Glioma Using an Integrated Meta-Clustering Framework

**DOI:** 10.64898/2026.08.27.747680

**Authors:** Bulidierxin Tuerhanbayi, Jieqiong Wang, Shibiao Wan

## Abstract

Pediatric low-grade glioma (pLGG) is the most common type of brain tumor in children, accounting for approximately 30% of all central nervous system tumors in children. pLGG has multiple molecular subtypes that differ in disease progression, recurrence patterns, and treatment responses. Conventional wet lab approaches including molecular profiling and histopathological studies for pLGG characterization are time consuming, costly, and laborious. Recently, methods based on artificial intelligence (AI) or machine learning (ML) have been widely used for pLGG molecular categorization, but most of them can only identify two or three pLGG subtypes. To more comprehensively characterize the molecular subtypes of pLGG and their potential biological and therapeutic significance, we develop an integrated meta-clustering approach, namely Meta-pLGG, that can explore high resolution molecular subtypes and their transcriptional heterogeneity for pLGG. Specifically, we first performed multiple rounds of random projection (RP) to generate dimension-reduced feature vectors from pLGG transcriptomics data, each of which was subsequently clustered by different clustering algorithms including hierarchical clustering, *K*-means, Self-Organizing Maps (SOM), Non-negative Matrix Factorization (NMF), Gaussian Mixture Model (GMM), and Spectral Clustering, as base clustering methods. Then, to yield robust clustering performance, we integrated the clustering results of these RP based individual clustering algorithms by adopting a weighted meta-clustering (wMetaC) approach. Results based on 532 pLGG patients suggested that our proposed approach demonstrated superior stability and discriminative powers for higher resolution pLGG subtyping compared to conventional approaches. Based on consensus matrix analysis, we identified two major pLGG mega-subtypes, with one further subdivided into three subgroups and the other into two. Then, we performed cluster specific differential gene expression analysis, molecular pathway analysis, and gene-drug-disease association analysis. The results showed that the identified five subgroups exhibited significant subtype-specific transcriptomic heterogeneity. In summary, our meta-clustering approach demonstrated much higher performance and robustness in identifying higher resolution molecular subtypes of pLGG, revealing the molecular heterogeneity within pLGG and potentially providing new insights for more precise molecular subtyping and precision therapy.

## Introduction

Pediatric low-grade glioma (pLGG) is one of the most common central nervous system tumors in children, accounting for approximately 30% of pediatric CNS tumors [1], and is classified as WHO grades 1-2 [2]. Although most patients with pLGG have a high overall survival rate, the disease remains clinically heterogeneous [3, 4]. These tumors often arise in functionally critical regions, such as the optic pathway, brainstem, and thalamus. As a result, complete surgical resection can be challenging in some patients, leading to long-term risks of recurrence and the need for multiple rounds of treatment [5].

Traditionally, pLGG classification relied on histopathological examination, which involves multiple steps including tissue processing, sectioning, staining, and microscopic evaluation by pathologists [6]. These procedures can be time consuming and labor intensive. In addition, tumors with similar histological features may have different molecular alterations, limiting the ability of morphology alone to fully characterize tumor heterogeneity [7]. Molecular and transcriptomic approaches can therefore provide additional information for more detailed tumor classification [8–10]. With the development of artificial intelligence (AI) and machine learning (ML) based approaches, unsupervised clustering has become an important strategy for identifying latent molecular subtypes [11, 12]. In glioma research, transcriptomic data have been increasingly used for molecular classification, as RNA sequencing (RNA-seq) enables genome wide characterization of gene expression patterns and facilitates the identification of molecularly distinct tumor groups [13].

Previous studies have shown that RNA-seq based clustering can reveal biologically meaningful subgroups within glioma. For example, Hernández et al. showed that transcriptomic profiles of pediatric astrocytoma revealed clear differences between low-grade and high-grade tumors [14]. Picard et al. identified two molecular subgroups of pilocytic astrocytoma through multi-omics analysis, characterized by distinct immune-response and neuronal-function features [15]. More recently, Kazerooni et al. integrated magnetic resonance imaging, machine learning, and RNA-seq data to classify pLGG into three clusters associated with immune activity and clinical outcomes [16]. Similarly, in adult lower-grade glioma, Lin et al. reported inflammation related RNA-seq subtypes with distinct immune microenvironment, mutational profiles, and survival outcomes [17]. Previous studies have mostly relied on a single clustering algorithm, making the results sensitive to the choice of method and potentially leading to inconsistencies and insufficient reproducibility in the classification results. The weighted meta-clustering framework integrates multiple base clustering solutions by incorporating weighting information to reduce the influence of less reliable clustering patterns and generate a more robust consensus classification [18–20].

To more comprehensively characterize the molecular heterogeneity of pLGG and explore its potential biological and therapeutic significance, we developed a meta-clustering method for pLGG subtype discovery, named Meta-pLGG. Meta-pLGG applies multiple rounds of RP to pLGG transcriptomic data, clusters each reduced representation using base clustering algorithms, and then integrates the clustering results using wMetaC to obtain robust consensus subtypes. By combining multiple data representations and clustering solutions, Meta-pLGG provides a systematic approach for pLGG subtype identification and may improve disease classification while offering insights into biologically and clinically relevant tumor subgroups.

## Results

### Meta-pLGG Framework Design

To more accurately and robustly identify molecular subtypes of pLGG, we developed an unsupervised clustering framework that integrates random projection, multiple base clutering methods and weighted meta-clustering (wMetaC), named Meta-pLGG. Rather than relying on a single clustering algorithm, Meta-pLGG intergrates clustering information generated from multiple low dimensional representations and clustering algorithms to reduce method specific variability and improve the stability of the subtype identification.

The Meta-pLGG architecture (**Fig. 1**) consists of three major components. First, the preprocessed RNA-seq expression matrix was used as the input for multiple rounds of random projection based dimensionality reduction. Through repeated RP, the high-dimensional gene expression matrix was mapped into multiple low-dimensional feature spaces, reducing the impact of noise and computational complexity. Second, within each low-dimensional space generated by random projection, six base clustering methods, including hierarchical [21], K-means [22], SOM [23], NMF [24], GMM [25], and Spectral [26] clustering, were independently applied to perform initial sample grouping. These methods characterize the similarity structure among samples from different perspectives and provide complementary clustering information. Third, all base clustering results generated across clustering methods were integrated using a weighted meta-clustering method called wMetaC. Instead of treating all clustering assignments equally, wMetaC quantifies the consistency of sample co-clustering first and uses this information to construct a weighted similarity between clusters. Clusters with similar sample compositions across multiple clustering solutions are subsequently grouped at the meta-clustering level, followed by a voting procedure to determine the final subtype assignment for each sample. This strategy enables Meta-pLGG to integrate consistent clustering patterns while reducing the influence of unstable individual clustering assignments. Ultimately, Meta-pLGG identified robust transcriptomic molecular subtypes of pLGG, which were further characterized through differential expression, pathway enrichment, cell-type enrichment, gene-drug-disease association, clinical information and survival analyses.

**Fig. 1.**
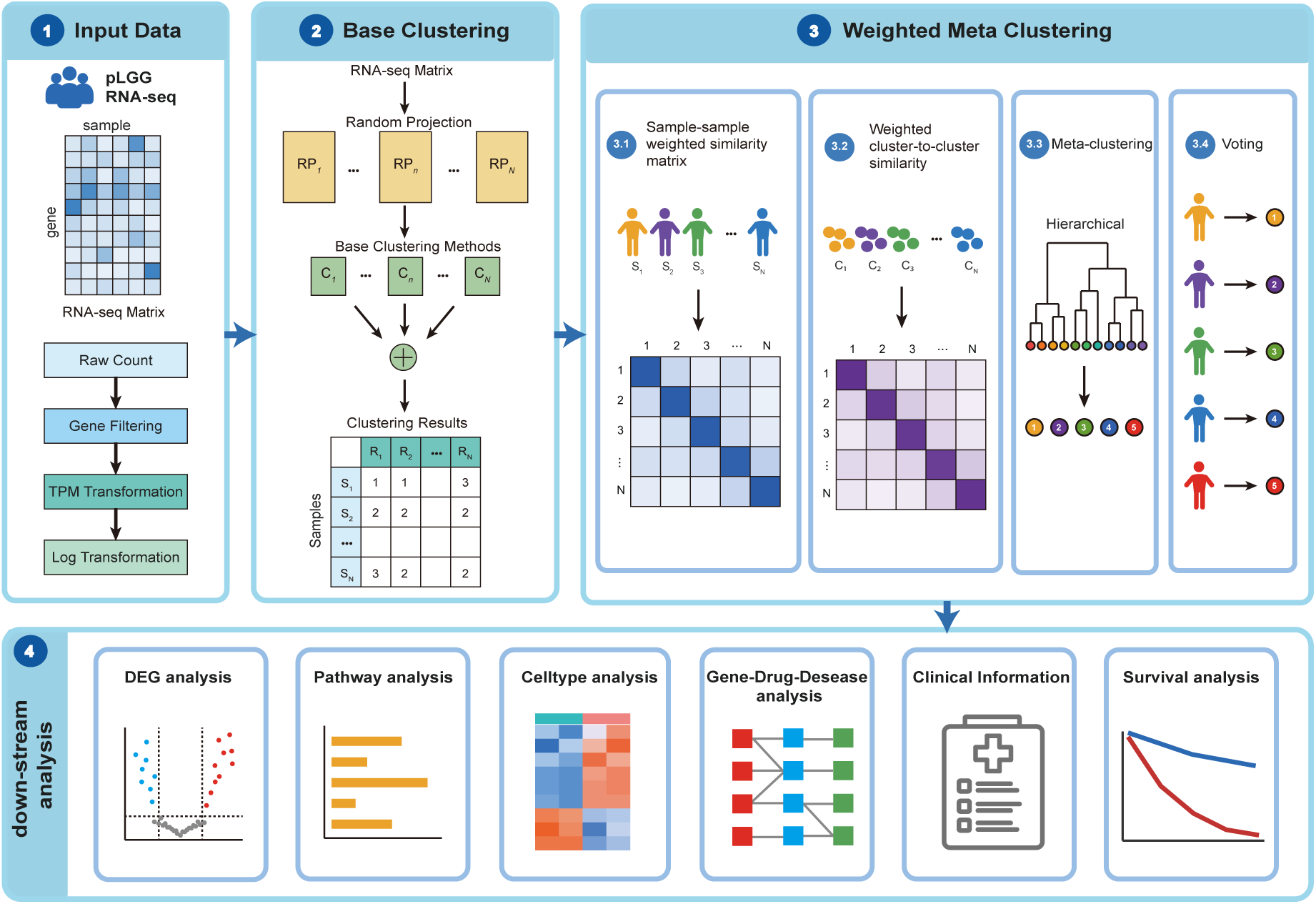
Design of Meta-pLGG. Meta-pLGG consists of these steps for pLGG subtype identification: RNA-seq data preprocessing, multiple random projection based dimensionality reduction, base clustering using hierarchical, K-means, SOM, NMF, GMM, and Spectral clustering, and weighted meta-clustering to generate final pLGG subtypes. RP: random projection, C: clustering methods. S: samples. R: clustering results. DEG: differentially expressed gene.

To determine the optimal combination of clustering methods, we systematically compared Meta-pLGG models constructed using the top six, top five, top four, top three, top two, and top one ranked clustering methods. Among these configurations, the model integrating all six clustering methods achieved the best performance (**Supplemental Figure 1**) and was therefore selected as the final Meta-pLGG architecture.

### Meta-pLGG Outperformed State-of-the-Art Clustering Methods for pLGG Subtyping

Meta-pLGG finally identified five robust transcriptomic clusters (**Fig. 2A**). We then further compared Meta-pLGG with six individual base clustering methods, including hierarchical, K-means, SOM, NMF, GMM, and Spectral clustering. Compared with these single base clustering methods, Meta-pLGG achieved a significantly higher silhouette index (**Fig. 2B**). These results indicate that by integrating multiple random projection spaces and multiple base clustering algorithms, Meta-pLGG can generate more robust and reliable transcriptomic subtype classifications of pLGG than baseline clustering methods.

**Fig. 2.**
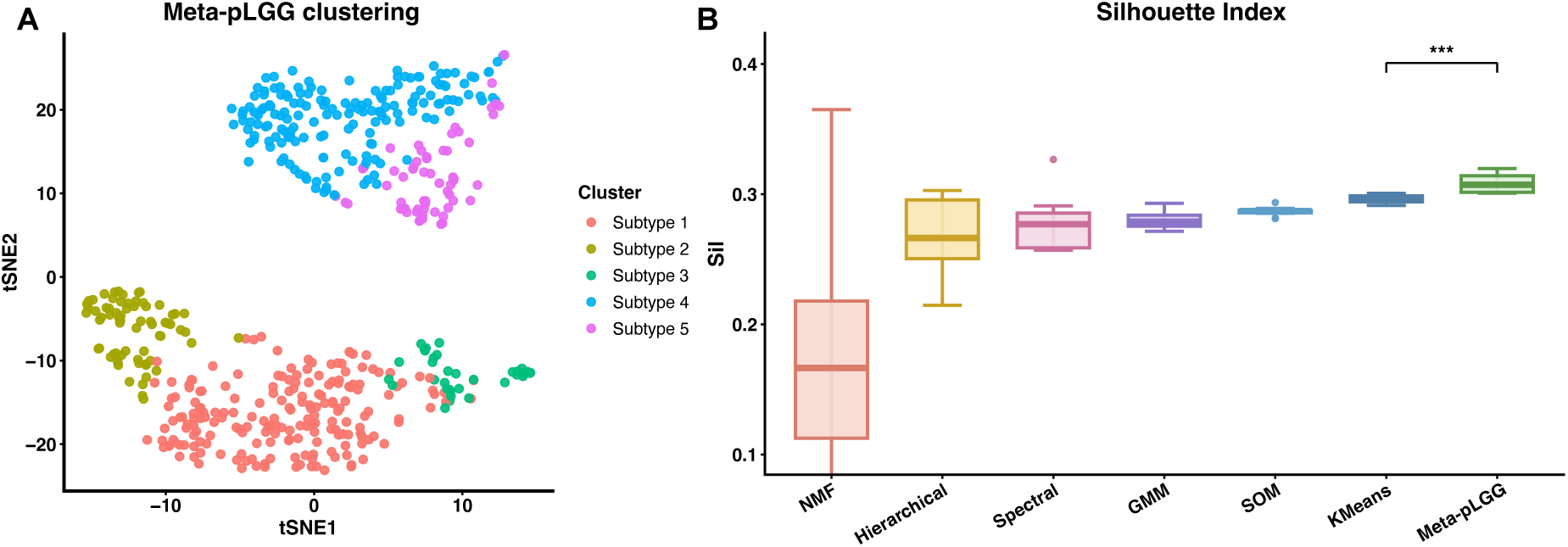
Meta-pLGG Outperformed SOTA Methods for Clustering pLGG. (**A**) t-SNE visualization of Meta-pLGG clustering results. Each dot represents one pLGG sample, and colors indicate the assigned subtype. Meta-pLGG identified five subtypes. (**B**) Meta-pLGG achieved higher silhouette index than the base clustering. **NMF:** non-negative matrix factorization; **GMM:**Gaussian mixture model; **SOM:** self-organizing map. *** represent *p* < 0.001.

### Subtype-Specific DGE Analysis Revealed Distinct Enriched Pathways Across Different pLGG Subtypes

To determine whether the five pLGG subtypes identified by Meta-pLGG exhibited distinct transcriptomic characteristics, we performed one-vs-rest differential gene expression analysis for each subgroup. The top 10 most representative subtype-specific genes from each subgroup were selected and visualized in a heatmap (**Fig. 3**), the five subgroups displayed clearly distinct gene expression patterns. These findings suggest that the Meta-pLGG subgroups are well separated at the transcriptomic level and may reflect underlying biological heterogeneity.

**Fig. 3.**
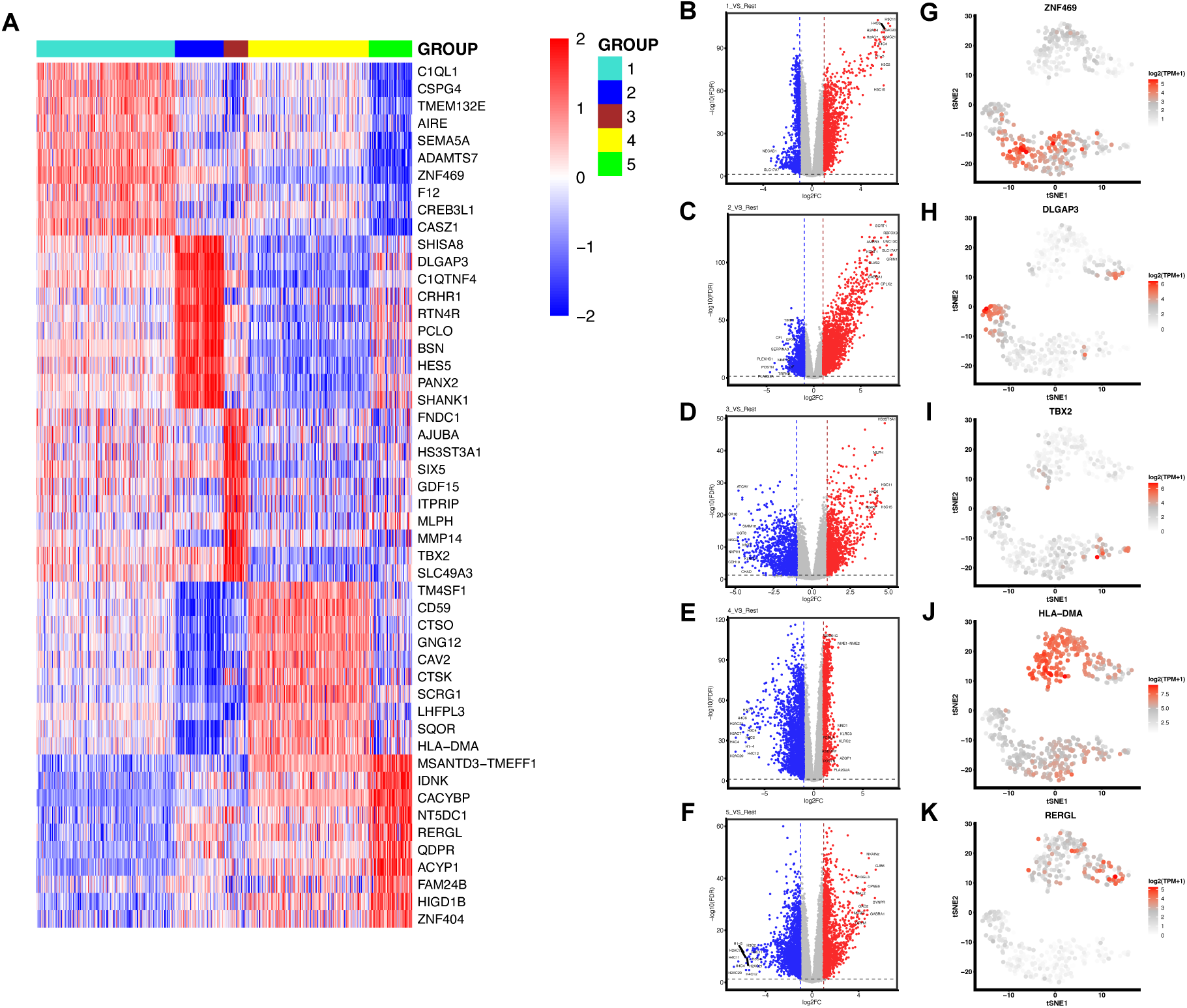
Subtype-Specific DGE Analysis Showed Distinct Expression Patterns. (**A**) Heatmap illustrating the top ten upregulated genes for each subtype in a one-vs-rest comparison. Volcano plots showing differentially expressed genes for (**B**) Subtype 1 vs rest. (**C**) Subtype 2 vs rest. (**D**) Subtype 3 vs rest. (**E**) Subtype 4 vs rest. (**F**) Subtype 5 vs rest. Significantly upregulated and downregulated genes were identified using thresholds of |logFC| > 1 and *p* value < 0.05. UMAP plots showing the relative expression of representative upregulated genes: (**G**) ZNF469 in Subtype 1, (**H**) DLGAP3 in Subtype 2, (**I**) TBX2 in Subtype 3, (**J**) HLA-DMA in Subtype 4, and (**K**) RERGL in Subtype 5. Higher expression levels are indicated by greater red intensity .

In Subtype 1, CSPG4 and ZNF469 showed relatively high expression. ZNF469 is a transcriptional regulator of collagen production and extracellular-matrix homeostasis [27]. Experimental depletion of ZNF469 has been shown to reduce collagen production and suppress ECM-associated transcriptional programs [28], whereas CSPG4 is a cell surface proteoglycan, which can regulates cancer cell migration, invasion, epithelial mesenchymal transition, and proliferation [29]. In Subtype 2, BSN and DLGAP3 showed subtype-specific expression. BSN encodes Bassoon, a core structural component of the presynaptic active-zone cytomatrix that organizes neurotransmitter-release sites and regulates synaptic-vesicle exocytosis [30].

DLGAP3 is a component of the postsynaptic density [31]. In Subtype 3, GDF15 and TBX2 were highly expressed. GDF15 is a member of the TGF-β superfamily and can be induced under cellular stress conditions. Previous studies have shown that elevated GDF15 levels are associated with pathological conditions such as inflammation and cancer, and that GDF15 may contribute to shaping the tumor microenvironment [32]. In glioblastoma models, GDF15 has also been reported to promote VEGFA expression and angiogenic communication between tumor cells and brain microvascular endothelial cells [33]. whereas TBX2 is a developmental transcription factor whose overexpression in glioblastoma promotes epithelial-to-mesenchymal-transition-like changes and increases cell migration and invasion [34]. In Subtype 4, HLA-DMA and CD59 were representative upregulated genes, consistent with the strong enrichment of antigen-processing and immune-response pathways. HLA-DMA encodes the α-chain of HLA-DM, a non-classical MHC class II heterodimer that promotes the selection and loading of stable peptide MHC class II complexes in antigen-presenting cells [35, 36], whereas CD59 prevents membrane-attack-complex formation by interacting with complement components C8 and C9, thereby protecting cells from complement-mediated damage [37]. In Subtype 5, ACYP1 and RERGL showed relatively high expression. ACYP1 encodes a cytosolic acylphosphatase involved in the hydrolysis of acylphosphates and has been linked to altered tumor-cell metabolism [38].

### GO Enrichment Analysis Revealed Distinct Enriched Pathways Across pLGG Subtypes

To further characterize the biological differences among the identified subtypes, we performed Gene Ontology (GO) enrichment analysis based on the upregulated genes in each subtype and compared the top 15 most significantly enriched GO terms. The results showed that each pLGG subtype was associated with distinct functional programs. Based on their dominant GO enrichment patterns and representative molecular features, the five subtypes were named as Extracellular Matrix (ECM)-α, Neuronal-α, ECM-β, Myeloid-antigen presentation (Myeloid-AP), and Neuronal-β subtypes, respectively.

In ECM-α (**Fig. 4B**), upregulated genes were mainly enriched in extracellular matrix related processes and components, including extracellular matrix organization, extracellular matrix structural constituents and mesenchyme development, indicating an extracellular matrix-rich and mesenchyme associated state. In Neuronal-α (**Fig. 4C**), upregulated genes were significantly enriched in neuronal synaptic structures and electrophysiological regulation, including neuron-to-neuron synapses, membrane potential regulation, and ion channel activity. These findings suggest that Neuronal exhibits a prominent neuronal and synaptic transcriptional program. In ECM-β (**Fig. 4D**), upregulated genes were enriched in extracellular matrix organization and collagen fibril organization, suggesting enhanced cell-matrix interactions. In Myeloid-AP (**Fig. 4E**), upregulated genes were strongly enriched in MHC class II protein complex assembly, peptidase inhibitor activity, and cell killing, indicating an immune-enriched state characterized by active antigen processing and presentation. Neuronal- β (**Fig. 5F**) was predominantly enriched in neurotransmitter secretion and transport and synaptic vesicles, indicating a neuronal secretory and synaptic signaling state.

**Fig. 4.**
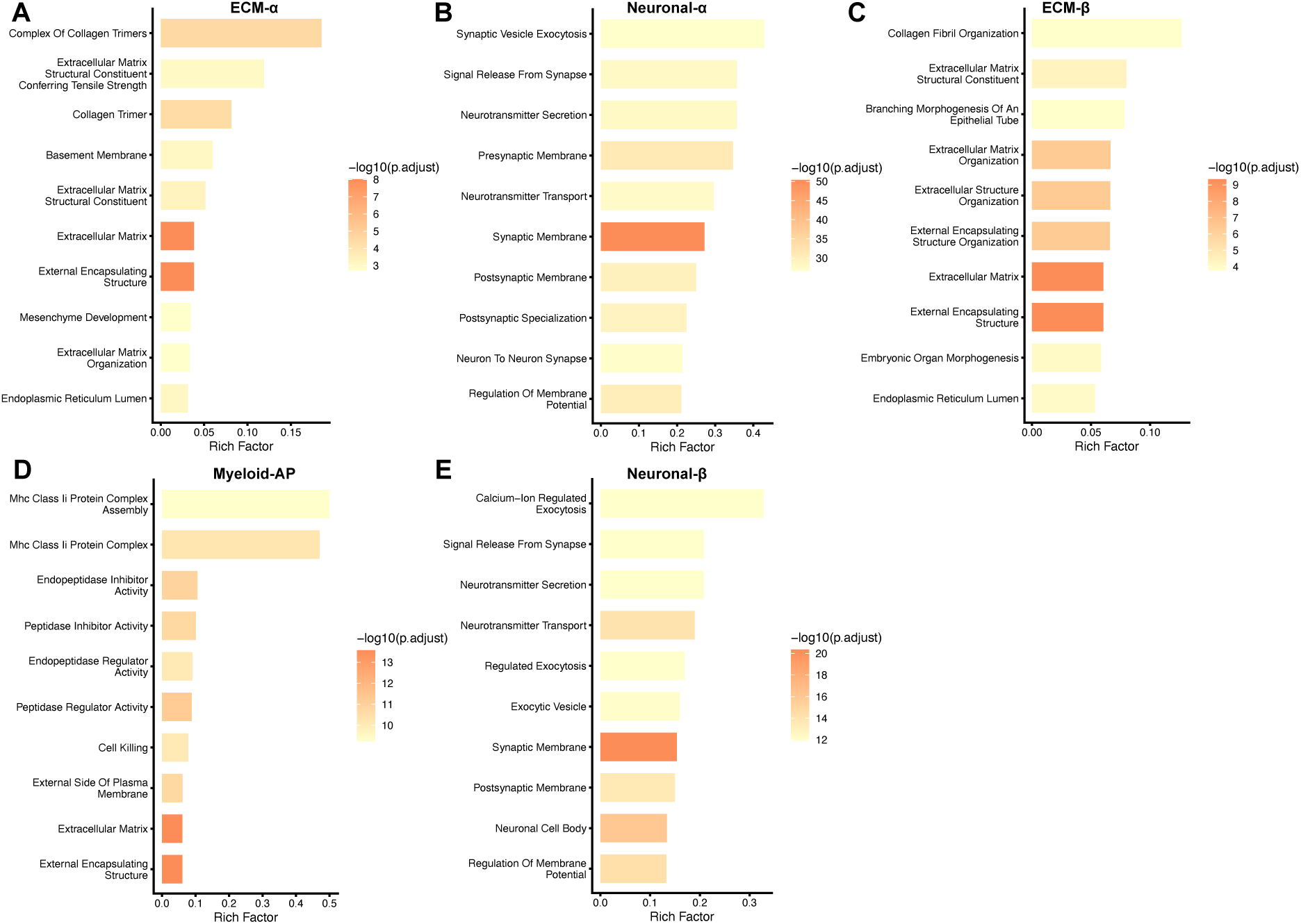
Pathway Analysis Revealed Functional Differences Across Subtypes. Bar plots showing the 10 most significant GO pathways enriched among the upregulated differentially expressed genes for: (**A**) Subtype ECM-α, (**B**) Subtype Neuronal-α, (**C**) Subtype ECM-β, (**D**) Subtype Myeloid-AP and (**E**) Subtype Neuronal-β. The Top 10 GO pathways are selected by adjusted *p* value and ranked by rich factor. ECM: extracellular matrix. AP: antigen presentation.

**Fig. 5.**
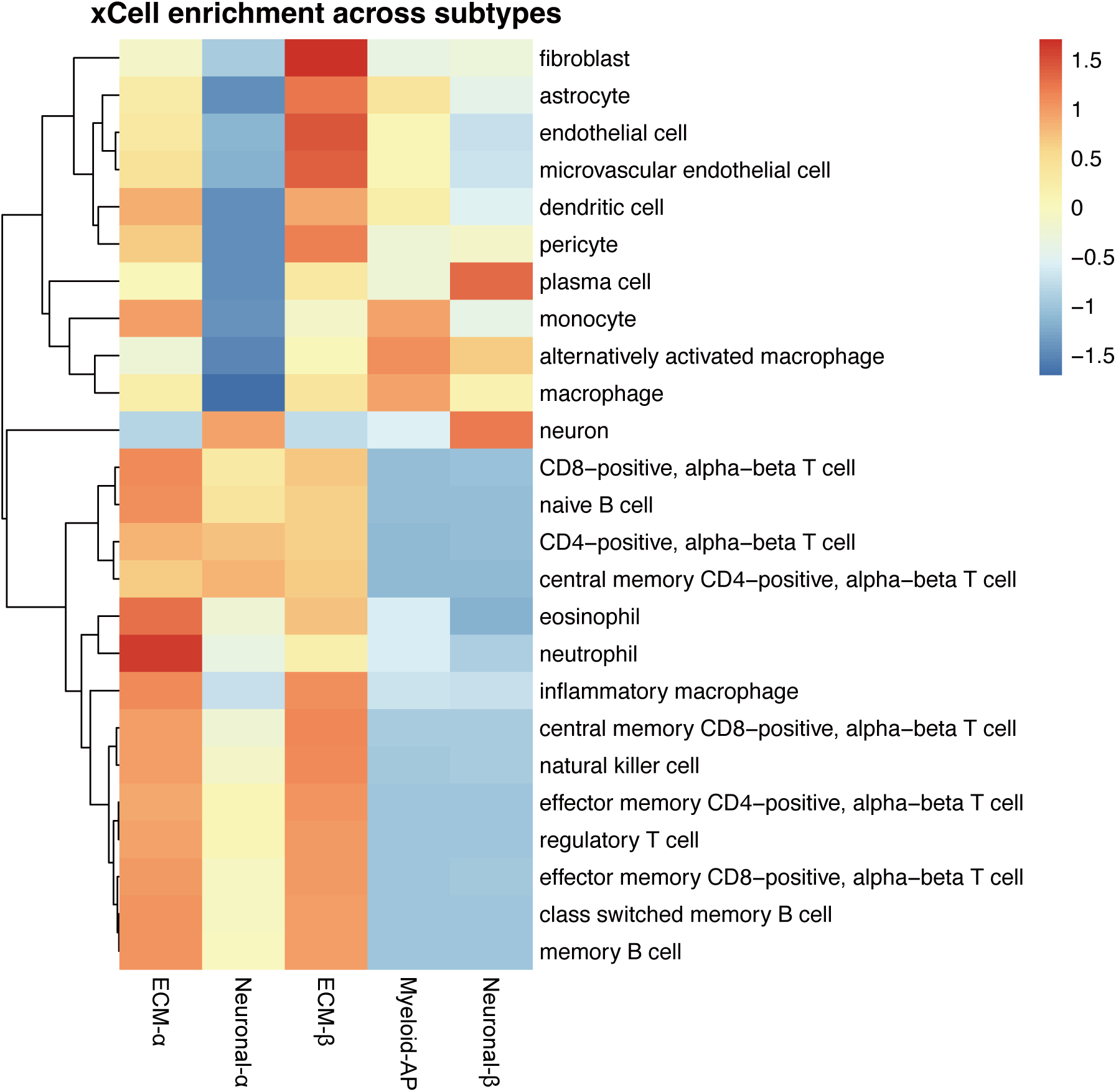
Heatmap of immune and stromal cell enrichment across the five subtypes based on xCell2 analysis. Differences in enrichment scores among the five subtypes (ECM-α, Neuronal- α, ECM-β, Myeloid-AP, and Neuronal-β) were evaluated using the Kruskal-Wallis test, with Benjamini-Hochberg correction for multiple comparisons. Cell types with a false discovery rate (FDR) < 0.05 were considered significantly different across subtypes, and representative cell types relevant to the glioma tumor microenvironment were selected for visualization. For each cell type, the mean xCell2 enrichment score was calculated within each subtype and standardized across the five subtypes using z-score transformation. Columns represent individual different subtypes, and rows represent immune and stromal cell types. ECM: extracellular matrix. AP: antigen presentation

### Immune cell enrichment analysis revealed distinct immune profiles across the pLGG subtypes

Cell type enrichment analysis revealed distinct cellular composition patterns across the five glioma subtypes. Cell-type enrichment was estimated from bulk RNA-seq data using xCell2. (**Fig. 5**). Subtypes ECM-α and ECM-β generally showed broader lymphocyte associated signals, including multiple T-cell and B-cell subsets. In contrast, most T-cell, B-cell, and natural killer-cell signatures were lower in Subtypes Myeloid-AP and Neuronal-β. ECM-α showed relatively high CD4⁺ and CD8⁺ T-cell, B-cell, natural killer-cell, neutrophil, and inflammatory macrophage signatures. ECM-β also exhibited relatively high memory T-cell, natural killer-cell, inflammatory macrophage, and dendritic-cell signatures. In addition, endothelial, microvascular endothelial, and fibroblast related signals were elevated in ECM-β. Neuronal-α showed a relatively high neuronal signature, whereas its myeloid, vascular, and stromal-cell-related signals were generally lower. Myeloid-AP displayed a distinct immune pattern. Monocyte, macrophage, and alternatively activated macrophage signatures were relatively high, whereas multiple T-cell, B-cell, and natural killer-cell signatures were low. Neuronal-β generally showed low mature T-cell, B-cell, natural killer-cell, inflammatory macrophage, and dendritic-cell signals, indicating a relatively immune-low microenvironment.

### Gene-Drug-Disease Association Analysis Identified Potential Subtype-Specific Target Genes and Drugs for pLGG

Then, we performed gene-drug-disease association analysis based on the upregulated differentially expressed genes in each subtype. Sankey plots were used to visualize potential relationships among genes, drugs, and diseases (**Fig. 6**).

**Fig. 6.**
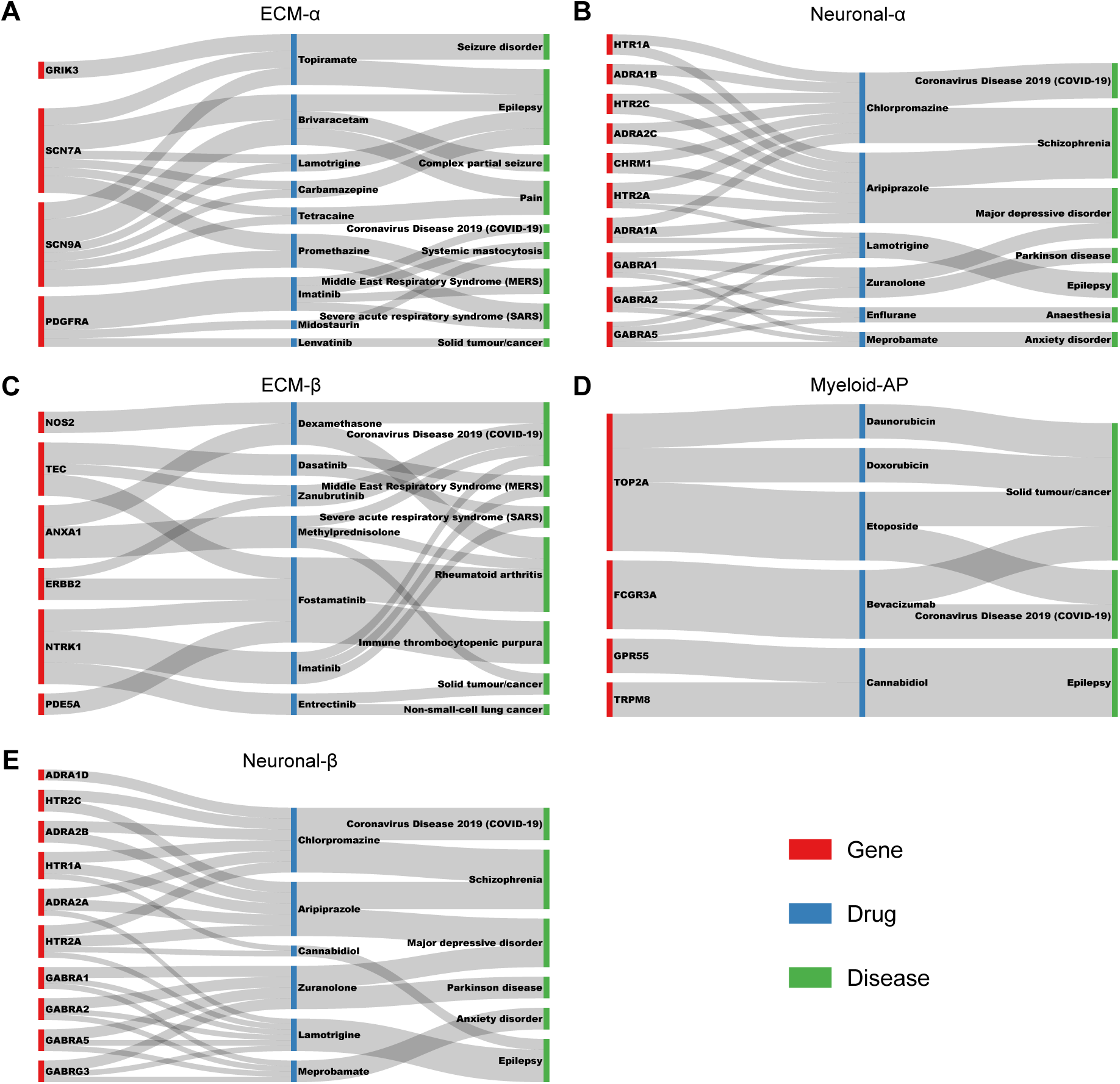
Gene-Drug-Disease Associations Suggested Distinct Therapeutic Relevance. Sankey diagrams illustrating the relationships among subtype-specific highly upregulated genes, associated drug and related diseases identified from the DrugBank database and Therapeutic Target Database (TTD) database. Results are shown for: (**A**) ECM-α, (**B**) Neuronal-α, (**C**) ECM-β, (**D**) Myeloid-AP and (**E**) Neuronal-β. ECM: extracellular matrix. AP: antigen presentation

In ECM-α (**Fig. 6A**), neuronal excitability-related genes, including SCN7A, and SCN9A, were linked to drugs such as brivaracetam and topiramate, with associated indications including epilepsy and pain. In addition, PDGFRA was connected to tyrosine kinase inhibitors, including imatinib and lenvatinib, with associated indications including systemic mastocytosis and solid tumors. In Neuronal-α (**Fig. 6B**), most upregulated genes were associated with chlorpromazine and aripiprazole, which are drugs commonly related to neuropsychiatric treatment, including antipsychotic and antidepressant related indications. In ECM-β (**Fig. 6C**), NTRK1 and ERBB2 were linked to targeted drugs, including entrectinib and imatinib, with associated indications such as solid tumors and non-small-cell lung cancer. In Myeloid-AP (**Fig. 6D**), FCGR3A was associated with bevacizumab, whereas TOP2A was linked to daunorubicin, doxorubicin, and etoposide, with associated indications including cancer. In Neuronal-β (**Fig. 6E**), upregulated genes were also mainly associated with chlorpromazine and aripiprazole. In addition, GABRA1 was associated with meprobamate and zuranolone, with related indications including anxiety disorder and epilepsy.

### Distinct Clinical and Molecular Features Across the Five pLGG Subtypes

The anatomical distribution of CNS regions differed across the five subtypes (**Fig. 7A**). Subtypes ECM-α and Myeloid-AP were mainly associated with posterior fossa tumors. Neuronal-α was primarily composed of hemispheric and posterior fossa tumors. ECM-β showed a relatively high proportion of ventricular tumors, whereas Neuronal-β was mainly associated with hemispheric tumors and included a comparatively large proportion of tumors involving mixed regions.

**Fig. 7.**
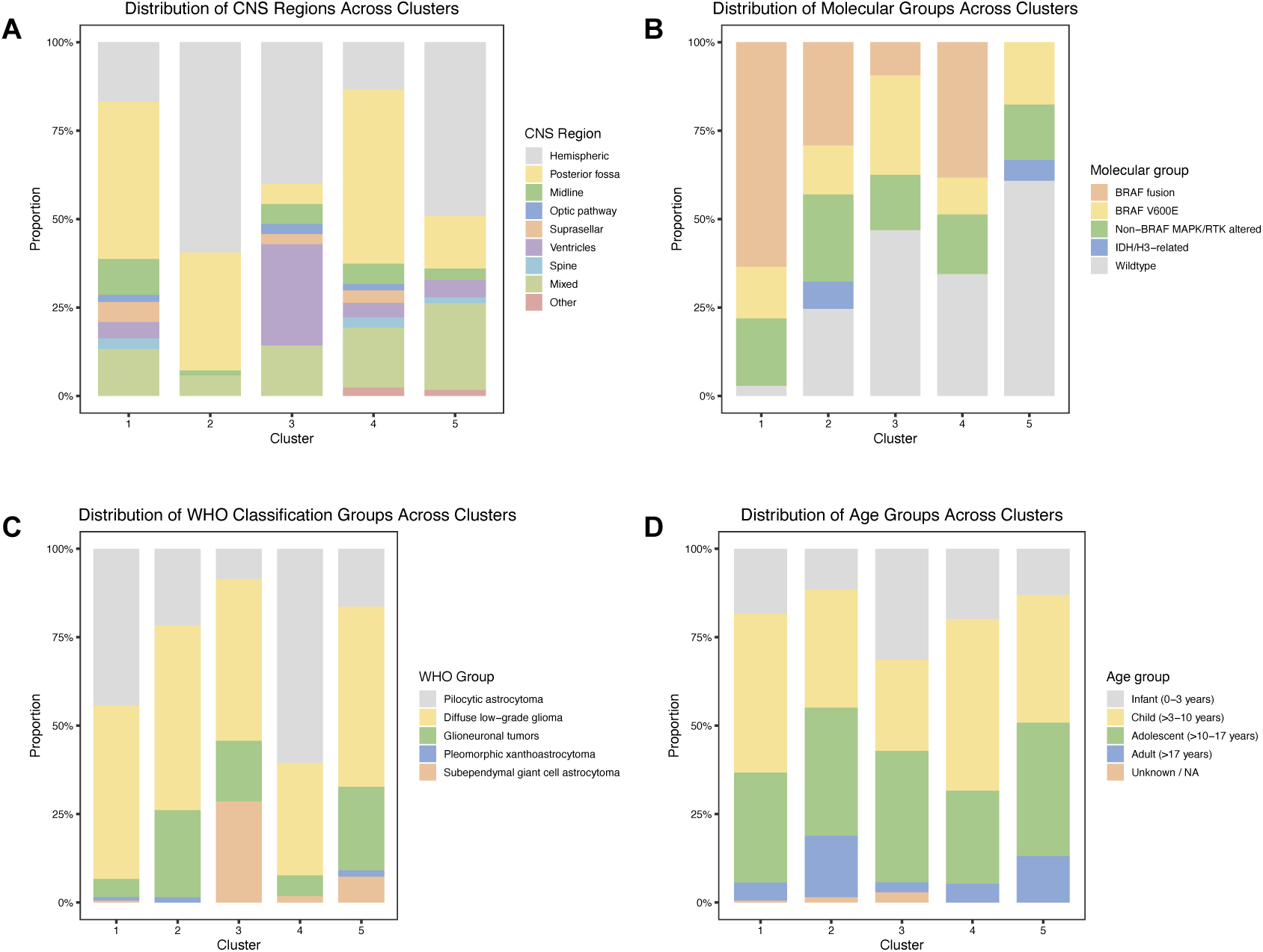
Clinical and molecular characteristics of the five Meta-pLGG subtypes. (**A**) Distribution of central nervous system (CNS) tumor locations across the five subtypes. (**B**) Distribution of molecular groups among the five subtypes. (**C**) Distribution of World Health Organization (WHO) diagnostic groups across the five subtypes. (**D**) Distribution of age groups among the five subtypes. Bar heights represent the proportion of patients within each category for the corresponding subtype. 1: ECM-α. 2: Neuronal-α 3: ECM-β. 4: Myeloid-AP. 5: Neuronal-β.

The distribution of molecular groups also varied across the five subtypes (**Fig. 7B**). ECM-α was predominantly characterized by BRAF fusion. Neuronal-α showed a mixed composition involving BRAF fusion, non-BRAF MAPK/RTK alterations, and wildtype tumors. ECM-β was mainly composed of wildtype tumors and also contained a relatively high proportion of BRAF V600E tumors. Myeloid-AP included substantial proportions of both BRAF fusion and wildtype tumors. In contrast, whereas Neuronal-β was predominantly composed of wildtype tumors and included a small proportion of IDH/H3-related gliomas.

The distribution of WHO classification groups further differed across the five Meta-pLGG subtypes (**Fig. 7C**). Subtypes ECM-α and Myeloid-AP were mainly composed of pilocytic astrocytoma and diffuse low-grade glioma. Neuronal-α was primarily characterized by diffuse low-grade glioma and glioneuronal tumors. In addition to diffuse low-grade glioma and glioneuronal tumors, ECM-β contained a relatively high proportion of subependymal giant cell astrocytoma. Neuronal-β was mainly composed of diffuse low-grade glioma and glioneuronal tumors, with smaller proportions of pilocytic astrocytoma and subependymal giant cell astrocytoma. The age distribution was similar across the five Meta-pLGG subtypes (**Fig. 7D**).

### Survival analysis revealed significant prognostic differences among the pLGG subtypes

Survival analysis showed that the five pLGG subtypes identified by Meta-pLGG had significantly different survival outcomes (**Fig. 8**). Subtypes Myeloid-AP and Neuronal-β showed relatively poor survival outcomes, whereas Subtypes ECM-α, ECM-β, and Neuronal- α had relatively favorable survival outcomes. Pairwise comparisons further supported this risk stratification. Myeloid-AP showed significant survival differences compared with Subtypes ECM-α, ECM-β, and Neuronal-α (**Supplemental table 1**). In addition, Neuronal-α and Neuronal-β also showed significant survival difference (**Supplemental table 1**).

**Fig. 8.**
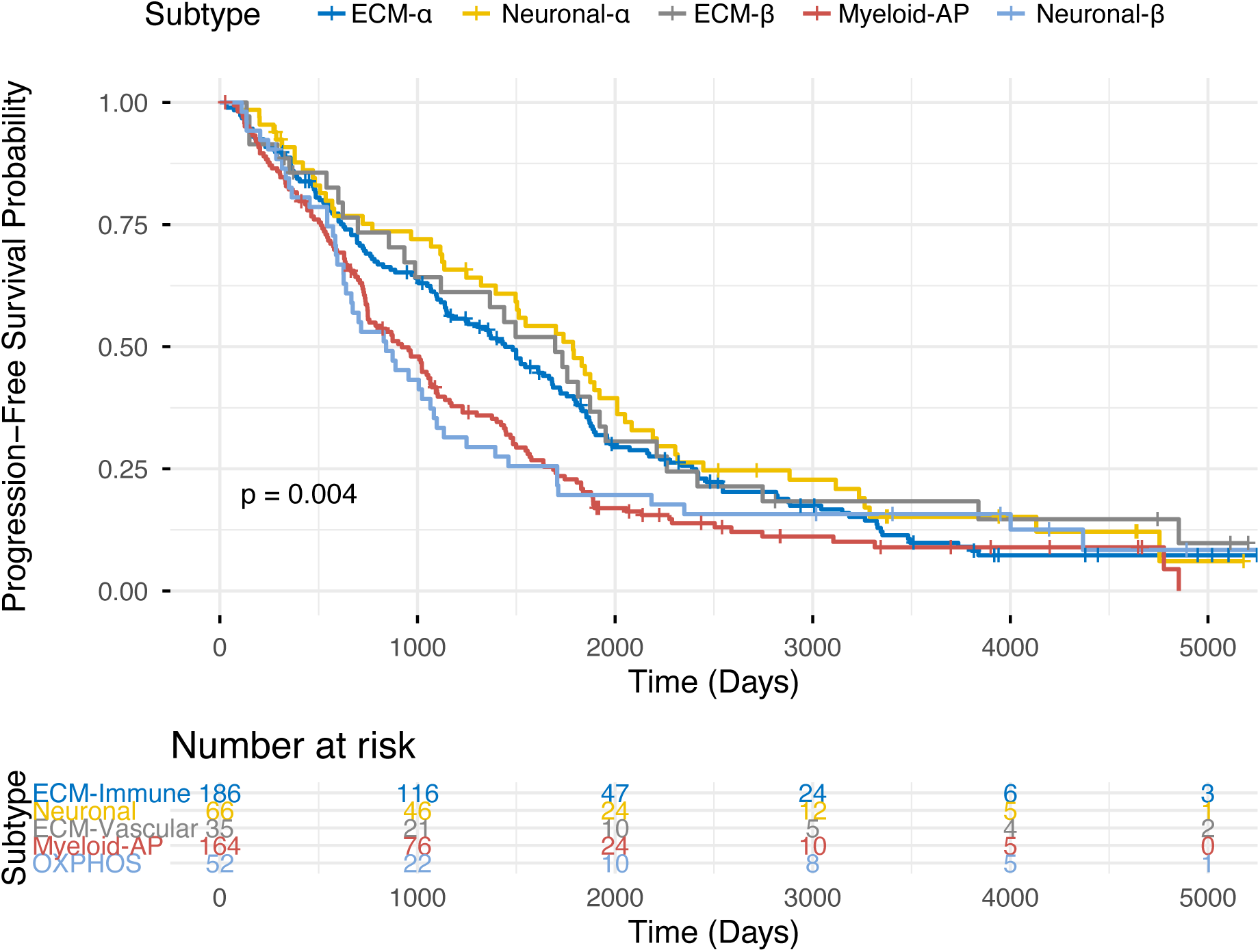
Survival Analysis Showed Subtype-Prognostic Differences. Kaplan-Meier survival curves and risk table for 5 subtypes identified by Meta-pLGG. *p* value < 0.05 was considered statistically significant.

### Discussion

Based on pLGG transcriptome expression data, we constructed a Meta-pLGG subtyping framework that integrates multiple random projections, various base clustering methods, and weighted meta-clustering. We further combined differential gene expression analysis, GO pathway enrichment analysis, immunce cell analysis, gene-drug-disease association analysis, clinical information analysis and survival analysis to systematically interpret the biological characteristics of the identified pLGG subgroups. Overall, our results suggest that Meta-pLGG not only improves clustering stability and subtype classification performance, but also identifies pLGG subgroups with distinct molecular features, potential drug associations, and different clinical outcomes.

First, compared with individual base clustering methods, Meta-pLGG showed better clustering performance and stability. A single clustering algorithm is often sensitive to parameter settings, initial values, and data noise. Therefore, different base clustering methods may generate inconsistent classifications for the same set of samples. In contrast, Meta-pLGG uses a weighted meta-clustering strategy to integrate multiple base clustering results. Stable sample relationships that repeatedly appear across different random projections and base clustering methods are assigned higher weights in the final classification. This strategy reduces the noise introduced by any single method and enables the identification of more robust molecular subtypes.

In addition, we found that Meta-pLGG achieved the highest silhouette index when more random projections were used. This finding suggests that multiple random projections can effectively reduce the randomness introduced by a single dimensionality reduction process, better preserve the structural information among samples, and improve the stability and separation of the final clustering results. Therefore, by combining multiple random projections, multiple base clustering algorithms, and weighted meta-clustering, Meta-pLGG provides a more reliable and robust analytical framework for transcriptomic classification of pLGG.

Differential expression analysis further supported the molecular distinction among the five Meta-pLGG subtypes. Each subtype exhibited a distinct set of upregulated genes, and several representative genes were consistent with the biological programs identified by pathway enrichment analysis. The ECM-α subtype was characterized by enrichment of extracellular matrix organization and relatively broad immune-cell enrichment. These findings suggest that the ECM-α subtype may represent a tumor microenvironment characterized by active extracellular matrix remodeling accompanied by immune involvement. Because extracellular matrix components can regulate immune-cell migration, retention, and cell-cell interactions [39], the coordinated enrichment of ECM and immune features may reflect close interactions between tumor cells and the surrounding microenvironment. Compared with ECM-α, the ECM-β subtype was characterized more strongly by stromal and vascular associated features, as reflected by the relatively high enrichment of fibroblasts, endothelial cells, microvascular endothelial cells. The Neuronal-α subtype showed prominent enrichment of neuron-related functions. Compared with Neuronal-α, the Neuronal-β subtype exhibited additional features related to oxidative phosphorylation. The relatively poor outcome of the Neuronal-β subtype may be associated with its enhanced mitochondrial respiratory and energy metabolism programs. Increased oxidative phosphorylation may support tumor cells survival [40, 41]. The comparatively lower enrichment of several lymphoid cell signatures in this subtype may also indicate a less active antitumor immune environment. The Myeloid-AP subtype exhibited prominent myeloid cell associated features and enrichment of antigen processing and presentation pathways, particularly MHC class II related processes. Tumor associated macrophages can adopt diverse functional states, and some myeloid populations may promote glioma growth by supporting tissue remodeling and immune suppression [42]. Therefore, the myeloid and antigen presentation signatures observed in this subtype may represent an immune active but not necessarily tumor restrictive microenvironment.

Gene-drug-disease association analysis further indicated distinct pharmacological backgrounds across the Meta-pLGG subtypes. ECM-α showed associations with neuronal excitability-related genes and PDGFRA linked tyrosine kinase inhibitors. ECM-β was mainly associated with targeted and anti-inflammatory treatments. Neuronal-α and Neuronal-β showed similar associations with neuroactive drugs, largely through neurotransmitter receptor genes, supporting the involvement of synaptic and neuromodulatory signaling. Myeloid-AP showed FCGR-related links to immune- and cancer associated drugs, consistent with its myeloid and antigen presentation features. These associations provide potential subtype-specific therapeutic hypotheses for further investigation.

The clinical and molecular characteristics further supported the biological heterogeneity of the five Meta-pLGG subtypes. Although ECM-α and ECM-β were both characterized by ECM-related transcriptomic pathways, their clinical and molecular profiles were distinct. ECM-α was predominantly associated with BRAF fusion and was more frequently located in the posterior fossa. In contrast, ECM-β contained a substantially larger proportion of wild-type tumors and fewer BRAF fusion-positive cases, with most tumors located in the hemispheric and ventricular regions. WHO classification also differed between these two subtypes. ECM-α consisted mainly of pilocytic astrocytoma and diffuse low-grade glioma, whereas ECM-β showed a higher proportion of glioneuronal tumors and subependymal giant cell astrocytoma.

This study also has some limitations. Meta-pLGG classification was mainly based on transcriptomic data, future studies incorporating multi-omics data may further improve the biological interpretation and clinical applicability of this classification framework. In addition, further experimental and clinical validation is needed to confirm the biological and therapeutic relevance of these findings.

## Methods

### Data Preprocessing

First, raw RNA-seq read counts were normalized to Transcripts Per Million (TPM). TPM normalization accounts for both gene length and differences in sequencing depth across samples, allowing gene expression levels to be more reliably compared across samples. Next, genes with low expression levels were filtered out to reduce background noise. Finally, the remaining TPM values were transformed using log_2_(TPM+1) to reduce the right-skewness of the expression distribution and the influence of extremely highly expressed genes.

### Random Projection

Random projection is a commonly used dimensionality reduction method [43, 44]. For data points in a high-dimensional space, random linear mapping can project the data into a lower-dimensional space while approximately preserving pairwise distances within an acceptable error range. Therefore, random projection can reduce computational complexity while retaining the major geometric structure of the original data [45]. Specifically, the original expression matrix contains *N* samples and *D* gene features, where each sample is represented by a *D*-dimensional expression vector. Random projection maps the original data from the *D*- dimensional space to a lower-dimensional subspace by multiplying the expression matrix with a randomly generated projection matrix, where *d* ≪ *D*. After random projection, each sample obtains a low-dimensional representation that approximately preserves the distance relationships among samples. In this study, the log2(TPM+1) transformed expression matrix was used for random projection analysis. To reduce the potential randomness introduced by a single random projection, multiple random projections were performed to generate multiple low-dimensional expression matrices.

### Weighted Meta Clustering

To integrate the clustering results generated from different random projections and base clustering algorithms, we applied an intergrated meta clustering method called weighted meta-clustering (wMetaC) [18]. The procedure consisted of five steps. First, for each pair of samples *i* and *j*, *s_i,j_* represents the probability that samples *i* and *j* are assigned to the same cluster. The pairwise weight was then calculated as:

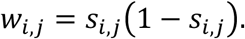

Second, the overall weight for sample *i* was calculated by summing its pairwise weights across all samples:

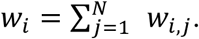

Third, all clusters generated across the base clustering results were pooled, and a weighted cluster-to-cluster similarity matrix was constructed. The similarity between two clusters *C_u_* and *C_v_* was calculated as:

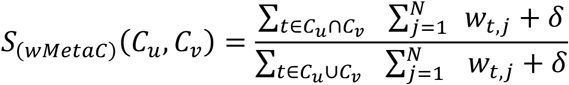

where *δ*=0.01. Fourth, hierarchical clustering using Ward’s method was applied to the weighted cluster similarity matrix to generate meta-clusters. Finally, a voting scheme was used to map the meta-cluster assignments back to individual samples, and the resulting consensus labels were used as the final Meta-pLGG subtype assignments.

### Subtype-Specific Differential Gene Expression Analysis

To identify subtype-specific transcriptomic features of pLGG, one-vs-rest differential gene expression analysis was performed for each subtype [46]. Genes with *p* value < 0.05 and log2 fold change > 1 were considered significantly upregulated subtype-specific genes in the corresponding subtype. Significantly upregulated genes from each one-vs-rest comparison were intersected with those identified in the corresponding pairwise subtype comparisons. Volcano plots were generated to visualize differentially expressed genes, with significance thresholds set as |log2FC| > 1 and adjusted *p* value < 0.05. Finally, tSNE visualization was performed based on the expression profiles of differentially expressed genes. Samples were colored according to the expression level of a representative significantly upregulated gene from each subtype.

### Pathway Enrichment Analysis

Pathway enrichment analysis was performed to characterize the biological functions associated with the differentially expressed genes identified for each subtype [47]. Subtype-specific upregulated genes were used as input for enrichment analysis. Enrichment analysis was conducted using the clusterProfiler R package with the enrichGO function, using the Gene Ontology (GO) annotation database. Biological Process, Cellular Component, and Molecular Function categories were evaluated, and terms with adjusted *p* values < 0.05 were considered significantly enriched. For visualization, the top 10 enriched terms were selected based on adjusted *p* values, with gene ratios indicating the proportion of subtype-specific genes associated with each term.

### xCell Immune Cells Analysis

To characterize relative differences in cellular enrichment across the five pLGG subtypes, xCell2 analysis was performed using the TPM expression matrix [48, 49]. xCell2 enrichment scores were calculated for each cell type in each sample based on predefined cell-type gene signatures. Differences in enrichment scores among the five subtypes were evaluated using the Kruskal-Wallis test, with Benjamini-Hochberg correction for multiple comparisons. Cell types with a false discovery rate (FDR) < 0.05 were considered significantly different across subtypes. For heatmap, the mean enrichment score of each cell type was calculated within each subtype and standardized across the five subtypes using z-score transformation.

### Gene-Drug-Disease Association Analysis

Drug related information was extracted from the DrugBank database [50], and target disease association information was obtained from the Therapeutic Target Database (TTD) [51]. Subtype-specific upregulated differentially expressed genes were matched to genes annotated in the integrated gene-drug-disease association table. Genes, drugs, and diseases were ranked according to their number of associated relationships, and the most frequently connected genes, drugs, and diseases were retained for visualization. The gene-drug-disease relationships were visualized using Sankey diagrams. Through this analysis, potential drug lists and their corresponding target genes were identified for each pLGG subtype.

### Kaplan-Meier Survival Curve Analysis

To evaluate prognostic differences among pLGG molecular subtypes, Kaplan-Meier survival analysis was performed using patients’ survival time and survival status. Patients were grouped according to their assigned molecular subtype, and survival curves were generated for each subtype. Differences in survival among groups were statistically evaluated using the log-rank test *p* < 0.05 was considered statistically significant. Patients who did not experience a death event by the end of follow up were treated as censored observations.

### Funding information

Research reported in this publication was supported by the U.S. National Science Foundation under Award Numbers 2500836 and 2614824, the National Cancer Institute of the National Institutes of Health under Award Number R03CA317707, the National Institute Of Alcohol Abuse And Alcoholism of the National Institutes of Health under Award Number R21AA032098, and the Office Of The Director, National Institutes Of Health of the National Institutes of Health under Award Number R03OD038391. This research was supported by the State of Nebraska through the Pediatric Cancer Research Group, part of the Child Health Research Institute. This work was also partially supported by the University of Nebraska Collaboration Initiative Grant from the Nebraska Research Initiative (NRI). The content is solely the responsibility of the authors and does not necessarily represent the official views of the funding organizations.

## Author contribution

The idea for this study was conceived of and designed by SW. SW and BT developed the method. BT performed the experiments and analyzed the data. All authors participated in the writing and revision of the paper. The manuscript was approved by all authors.

## Data Availability

All data used in this study are publicly available from a recent pLGG study (Kazerooni et al., Nature Communications, 2025).

## Code availability

All code used for Meta-pLGG is publicly available and can be found on GitHub at https://github.com/wan-mlab/Meta-pLGG.

