## Supplemental figures and tables for "High-Resolution Subtyping of Pediatric Low-Grade Glioma Using an Integrated Meta-Clustering Framework"

**Supplemental figure and table**

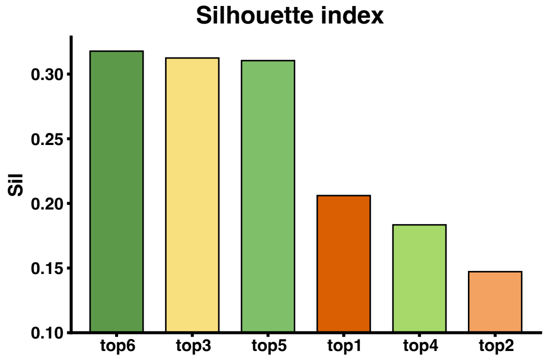


**Supplemental Figure 1. Comparison of weighted meta-clustering performance across different combinations of base clustering methods.** Weighted meta-clustering was performed using the top six, five, four, three, two, and one base clustering methods. Clustering performance was evaluated using the silhouette index. The combination incorporating all six base clustering methods achieved the best performance.

**Supplemental table 1 Pairwise Survival Analysis Result.** Pairwise survival comparisons were performed between Meta-pLGG subtypes using the log-rank test. The table shows the *p* values for each subtypes comparison, indicating whether survival outcomes differed significantly between two groups. *p* values < 0.05 were considered statistically significant.

| Group | *p*_value |
|---|---|
| ECM-α vs Neuronal-α | 0.24747 |
| ECM-α vs ECM-β | 0.52891 |
| ECM-α vs Myeloid-AP | 0.0038193 |
| ECM-α vs Neuronal-β | 0.18755 |
| Neuronal-α vs ECM-β | 0.86224 |
| Neuronal-α vs Myeloid-AP | 0.0017865 |
| Neuronal-α vs Neuronal-β | 0.044594 |
| ECM-β vs Myeloid-AP | 0.027942 |
| ECM-β vs Neuronal-β | 0.12202 |
| Myeloid-AP vs Neuronal-β | 0.82746 |
